# Meta-taxonomic Analysis of Prokaryotic and Eukaryotic Gut Flora in Stool Samples from Visceral Leishmaniasis Cases and Endemic Controls in Bihar State India

**DOI:** 10.1101/630624

**Authors:** Rachael Lappan, Cajsa Classon, Shashi Kumar, Om-Prakash Singh, Ricardo V. de Almeida, Jaya Chakravarty, Poonam Kumari, Sangeeta Kansal, Shyam Sundar, Jenefer M. Blackwell

**Affiliations:** Telethon Kids Institute, The University of Western Australia, Nedlands, Western Australia, Australia; Department of Microbiology, Tumor and Cell Biology, Karolinska Institutet, Solna, Sweden; Department of Medicine, Institute of Medical Sciences, Banaras Hindu University, Varanasi, India; Departamento de Bioquímica, Centro de Biociências, Universidade Federal do Rio Grande do Norte, Natal, Brazil; Department of Pathology, University of Cambridge, Cambridge, UK

**Author notes:** Corresponding Author: Professor Jenefer Blackwell, Telethon Kids Institute, PO Box 855, West Perth, Western Australia 6872, Australia. Equal senior authors (Shyam Sundar, Jenefer M. Blackwell).

## Abstract

Visceral leishmaniasis (VL) caused by *Leishmania donovani* remains of public health concern in rural India. Those at risk of VL are also at risk of other neglected tropical diseases (NTDs) including soil transmitted helminths. Intestinal helminths are potent regulators of host immune responses sometimes mediated through cross-talk with gut microbiota. We evaluate a meta-taxonomic approach to determine the composition of prokaryotic and eukaryotic gut microflora using amplicon-based sequencing of 16S ribosomal RNA (16S rRNA) and 18S rRNA gene regions. The most abundant bacterial taxa identified in faecal samples from Bihar State India were *Prevotella* (37.1%), *Faecalibacterium* (11.3%), *Escherichia-Shigella* (9.1%), *Alloprevotella* (4.5%), *Bacteroides* (4.1%), *Ruminococcaceae* UCG-002 (1.6%), and *Bifidobacterium* (1.5%). Eukaryotic taxa identified (excluding plant genera) included *Blastocystis* (57.9%; Order: Stramenopiles), *Dientamoeba* (12.1%; Family: Tritrichomonadea), *Pentatrichomonas* (10.1%; Family: Trichomonodea), *Entamoeba* (3.5%; Family: Entamoebida), Ascaridida (0.8%; Family: Chromodorea; concordant with *Ascaris* by microscopy), Rhabditida (0.8%; Family: Chromodorea; concordant with *Strongyloides*), and Cyclophyllidea (0.2%; Order: Eucestoda; concordant with *Hymenolepis*). Overall alpha (Shannon’s, Faith’s and Pielou’s indices) and beta (Bray-Curtis dissimilarity statistic; weighted UniFrac distances) diversity of taxa did not differ significantly by age, sex, geographic subdistrict, or VL case (N=23) *versus* endemic control (EC; N=23) status. However, taxon-specific associations occurred: (i) Ruminococcaceae UCG-014 and Gastranaerophilales_uncultured bacterium were enriched in EC compared to VL cases; (ii) *Pentatrichomonas* was more abundant in VL cases than in EC, whereas the reverse occurred for *Entamoeba*. Across the cohort, high *Escherichia-Shigella* was associated with reduced bacterial diversity, while high *Blastocystis* was associated with high bacterial diversity and low *Escherichia-Shigella*. Individuals with high *Blastocystis* had low Bacteroidaceae and high *Clostridiales* vadin BB60 whereas the reverse held true for low *Blastocystis*. This scoping study provides useful baseline data upon which to develop a broader analysis of pathogenic enteric microflora and their influence on gut microbial health and NTDs generally.

**Author Summary:** Visceral leishmaniasis (VL), also known as kala-azar, is a potentially fatal disease caused by intracellular parasites of the *Leishmania donovani* complex. VL is a serious public health problem in rural India, causing high morbidity and mortality, as well as major costs to local and national health budgets. People at risk of VL are also at risk of other neglected tropical diseases (NTDs) including soil transmitted helminths (worms). Intestinal worms are potent regulators of host immune responses often mediated through cross-talk with gut bacteria. Here we have used a modern DNA sequencing approach to determine the composition of microbiota in stool samples from VL cases and endemic controls. This allows us to determine all bacteria, as well as all single-celled and multicellular organisms, that comprise the microorganisms in the gut in a single sequencing experiment from a single stool sample. In addition to providing valuable information concerning commensal and pathogenic gut micro-organisms prevalent in this region of India, we find some specific associations between single-celled gut pathogens and VL case status.

## Introduction

Visceral leishmaniasis (VL), also known as kala-azar, is a potentially fatal disease caused by obligate intracellular parasites of the *Leishmania donovani* species complex. VL is a serious public health problem in indigenous and rural populations in India, causing high morbidity and mortality, as well as major costs to both local and national health budgets. The estimated annual global incidence of VL is 200,000 to 400,000, with up to 50,000 deaths annually occurring principally in India, Bangladesh, Sudan, South Sudan, Ethiopia and Brazil [1].

The outcome of infection with *L. donovani* depends on the host immune response [2], that in turn could be influenced by other factors, one of which could be co-infection with other pathogens. Often those at risk of VL are also at risk for other neglected tropical diseases (NTDs) such as infestation with soil transmitted helminths or infection with lymphatic filariasis or *Schistosoma mansoni*. Intestinal helminths are particularly potent regulators of their host’s immune response and can ameliorate inflammatory diseases [3]. Few published studies are available on interaction between *L. donovani* infection and other NTDs. Hassan and coworkers [4] showed that, despite the development of a functional anti-*L. donovani* T helper 1 response, mice with established *S. mansoni* infections fail to control *L. donovani* growth in the liver and spleen. Similarly, we recently demonstrated [5] that chronic infection with intestinal nematode *Heligmosomoides polygyrus* exacerbated secondary *L. donovani* infection in mice, resulting in higher parasite burdens in liver and spleen compared to worm free mice. This increased parasite load was accompanied by increased interleukin-4 and interleukin-10 transcription in spleens. In contrast, the presence of the filarial parasite *Brugia malayi* L3/adult worms inhibited progression of *L. donovani* infection associated with a T helper 1 response in a hamster model [6]. However, much may depend on the timing of infection with the other NTD and with *L. donovani*. A chronic helminth infection such as may be expected in areas endemic for VL may have a different effect compared to a newly established primary infection. In humans, the only co-infection studies reported to date are in relation to American tegumentary leishmaniasis [7–9]. In an initial study of 120 patients with cutaneous leishmaniasis caused by *L. braziliensis* [8] patients co-infected with helminths took longer to heal than helminth-free patients. However, in a follow up study of 90 patients there was no statistically significant benefit to treating patients with anti-helminthics [9]. In a more recent retrospective cohort of 109 L. braziliensis patients from Brazil [7] it was shown that patients with positive parasitological stool examinations had more mucosal lesions and more often had a poor response to therapy. Interestingly, these effects were observed for total intestinal helminths (based cumulatively on Ancylostomidae, *Ascaris lumbricoides*, *Strongyloides stercoralis*, *Trichuris trichuria*, *Enterobius vermicularis*, *S. mansoni*), and for *A. lumbricoides* alone or nematodes cumulatively (*A. lumbricoides*, *Str. stercoralis*, *T. trichuria, E. vermicularis*), but not for protozoan (based cumulatively on *Blastocystis hominis*, *Entamoeba coli*, *E. histolytica*, *Giardia lamblia*, *Endolimas nana*) infections.

One limiting factor for wide-ranging analysis of gut pathogen load has been the specialized and time-consuming processes involved in microscopic identification of parasites. While multiplex PCR for fecal DNA samples is facilitating larger scale studies for specific helminth and protozoan gut pathogens [10] this does not provide data on the potential influence of pathogenic species on gut microbes more broadly. For example, it was recently reported that the gut microbiome contributes to impaired immunity in pulmonary tuberculosis patients in India [11]. The presence of gut helminths is known to influence the gut microbiome [12, 13], and there is evidence [3] of important cross-talk between the gut microbiome and helminths in mediating host immunological effects. A meta-taxonomic approach to look at both commensal and pathogenic species in the gut would therefore be of great value. Numerous studies have used high-throughput, massively parallel amplicon-based sequencing of 16S ribosomal RNA (16S rRNA) gene regions to study the prokaryotic composition of the gut microflora [14–16], including studies in India [11, 17-21]. Recent studies in mammalian hosts [22] have also assessed parasite diversity using amplicon-based high-throughput sequencing of 18S ribosomal RNA (18S rRNA) gene regions and classified sequence reads into multiple parasite groups. Comparison of results with standard methods including microscopic observation of helminths in the intestines, and PCR amplification/ sequencing of 18S rDNA from isolated single worms, suggests that this new technique is reliable and useful to investigate eukaryotic parasite diversity in fecal samples. However, this approach has not to our knowledge been used in humans. Here we present a scoping study that uses 16S rRNA and 18S rRNA meta-taxonomic analysis of fecal samples with the primary aims of determining the composition of gut prokaryotic and eukaryotic microflora in an area of India endemic for VL and whether this differs between VL cases and non-VL endemic controls. We compare microscopic determination of helminth infections with 18S rRNA data, and report on secondary aims of determining the potential influence of prokaryotic or eukaryotic taxa known to contain pathogenic species on gut microbial diversity.

## Methods

### Study Subjects

Samples were collected between May and June 2017 at the Kala-azar Medical Research Center (KAMRC), Muzaffarpur, Bihar, India. Active VL cases (N=23) were diagnosed by experienced clinicians based on clinical signs, including fever (>2 weeks), splenomegaly, positive serology for recombinant antigen (r)-K39 and/or by microscopic demonstration of *Leishmania* amastigotes in splenic aspirate smears. Fecal samples (cf. below) were collected prior to treatment. Endemic controls (EC; N=23) that did not have VL were attendants accompanying VL patients at KAMRC (N=16) or from field locations nearby (N=7). Basic demographic details (age and sex) on participants are provided in Table 1. Metadata by individual are provided in S1 Table.

**Table 1.**
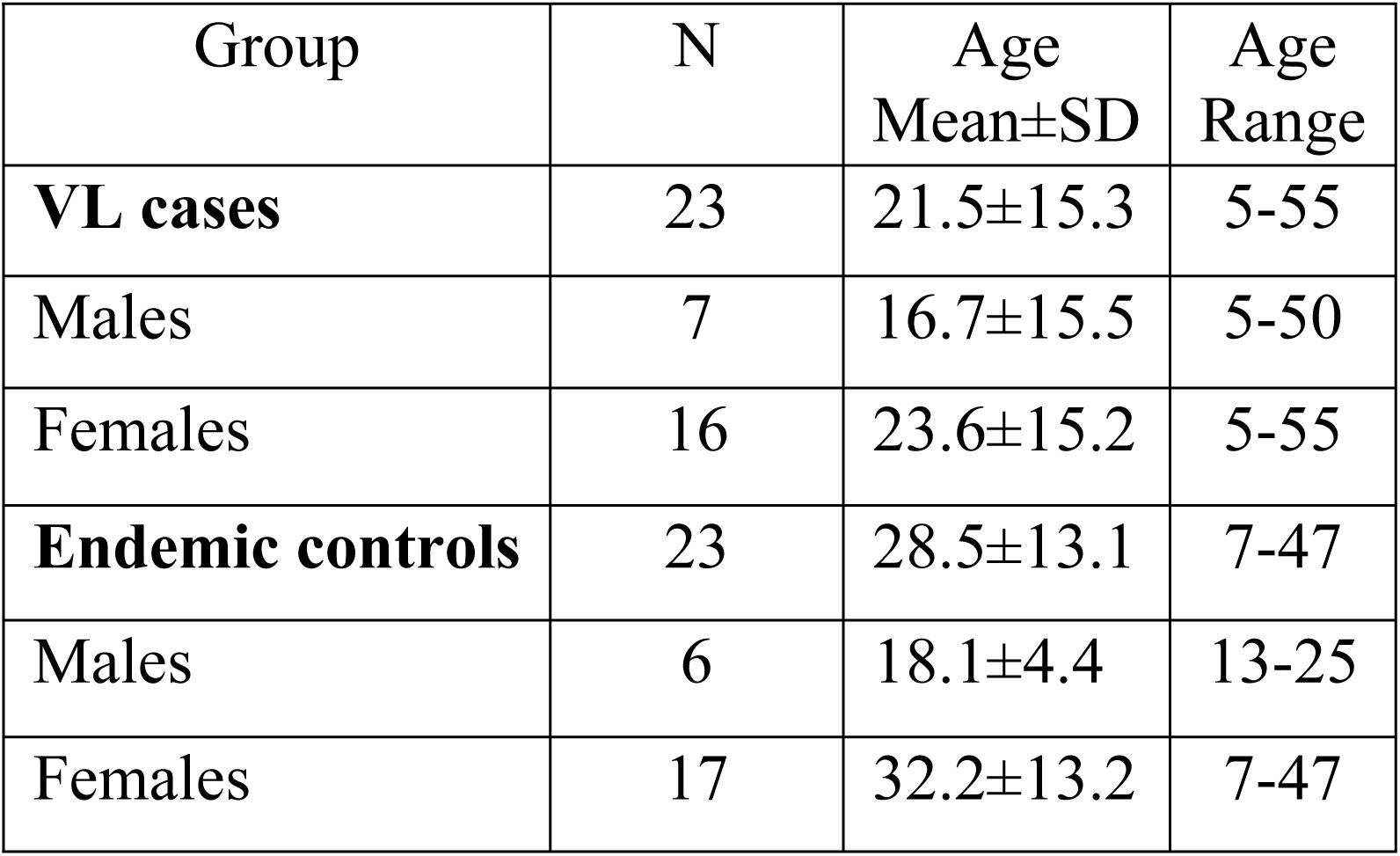
Basic demographic details of study participants. Full metadata are provided in S1 Table.

| Group | N | Age<br>Mean±SD | Age<br>Range |
| --- | --- | --- | --- |
| **VL cases** | 23 | 21.5±15.3 | 5-55 |
| Males | 7 | 16.7±15.5 | 5-50 |
| Females | 16 | 23.6±15.2 | 5-55 |
| **Endemic controls** | 23 | 28.5±13.1 | 7-47 |
| Males | 6 | 18.1±4.4 | 13-25 |
| Females | 17 | 32.2±13.2 | 7-47 |

### Ethics

The enrolment of human subjects complies with the principles laid down in the Helsinki declaration. Institutional ethical approval (reference numbers: Dean/2017/EC.148) was obtained from the ethical review board of Banaras Hindu University (BHU), Varanasi, India. Informed written consent was obtained from each participant at the time of enrolment, or from their legal guardian if they were under 18 years old. Only patients who had not previously received treatment and who agreed to participate in the study were enrolled. All clinical treatment and follow-up records were maintained using standardised case report forms on an electronic server.

### Stool collection

Stool collection containers and toilet accessories (to avoid urine contamination; OM-AC1; DNA Genotek) were distributed to the study participants and the containers collected the following morning. The sample was divided into two parts: 200 µg to OMNIgene GUT tubes (OM-200; DNA Genotek), and approximately 1 gram was concentrated by the formalin-ethyl acetate sedimentation method and used for diagnostic microscopy.

### Parasite microscopy

For microscopy, faeces were homogenized in 10 ml of 10% formalin containing 0.1% Triton-X-100 and kept at room temperature for at least 30 minutes for fixation. Samples were then filtered through two layers of gauge mesh, centrifuged 1500 *g* for 3 minutes, and the supernatant discarded. Seven ml of 10% formalin was added followed by 3 ml ethyl acetate, and samples shaken for 30 seconds and centrifuged 1500 *g* for 3 minutes. Supernatants were discarded and saline added to the sediment, which was then analysed under a light microscope at 10x magnification for the presence of helminth eggs.

### DNA extraction

A portion (200μg) of each stool sample (as above) was transferred to OMNIgene® GUT tubes (DNA Genotek) following the OMNIgene® GUT protocol for transferring specimens from the collection tubes to this kit (protocol PD-PR-00434). The specimen was homogenised and stabilised within these tubes prior to DNA extraction. Genomic DNA was extracted with the QIAamp PowerFecal DNA Kit (QIAGEN) following the manufacturer’s protocol with the exception that proteinase K was added after the bead beating step and the sample incubated at 65°C for 30 minutes. This was to ensure removal of histones from eukaryotic DNA which could inhibit downstream amplification. The remainder of the protocol was carried out as per manufacturer’s instructions. Purified genomic DNA was eluted in 100μl of PCR-grade water. Genomic DNA samples were shipped to the Telethon Kids Institute in Perth, Australia, at ambient temperature for further analysis and stored at - 20°C upon arrival.

### Sequencing controls

As a positive control for the 16S rRNA and 18S rRNA sequencing we used the ZymoBIOMICS Microbial Community DNA Standard (Integrated Sciences, Chatwood, NSW, Australia) which comprised DNA for 8 bacterial (*Listeria monocytogenes* - 12%, *Pseudomonas aeruginosa* - 12%, *Bacillus subtilis* - 12%, *Escherichia coli* - 12%, *Salmonella enterica* - 12%, *Lactobacillus fermentum* - 12%, *Enterococcus faecalis* - 12%, *Staphylococcus aureus* - 12%) and 2 eukaryotic (*Saccharomyces cerevisiae* - 2%, and *Cryptococcus neoformans* - 2%) species. To ensure that amplification of 16S rRNA and 18S rRNA amplicons worked well in the presence of human DNA, we included replicates of the positive control with human DNA spiked in at 0%, 0.1%, 0.5%, 1%, and 10% of the total 20 ng reaction, as indicated. For the 18S rRNA positive control we also spiked in DNA from two protozoan parasites *Leishmania major* (1.2 ng/20 ng reaction) and *Toxoplasma gondii* (1.2 ng/20 ng reaction). Negative controls included 4 unused OMNIgene® GUT tubes processed for DNA in the Indian laboratory using the QIAamp PowerFecal DNA Kit (QIAGEN) kits as for preparation of experimental sample DNAs. These were included in the preparation of amplicons for sequencing.

### Amplicon sequencing

Genomic DNA was quantified with the Qubit 2.0 dsDNA HS assay kit (Invitrogen) and diluted to 10 ng/µl for amplicon PCR. Aliquots for sequencing of 16S rRNA gene were sent to the Australian Genome Research Facility (AGRF) for amplification, Nextera XT dual-indexing, pooling and sequencing. The V3-V4 region of the gene was amplified with primers 341F (5’CCTAYGGGRBGCASCAG) and 806R (5’GGACTACNNGGGTATCTAAT) [23]. Aliquots for sequencing of the 18S rRNA gene were first amplified with the V9 region primers 1391f (5’**TCGTCGGCAGCGTCAGATGTGTATAAGAGACAG**GTACACACCGCCCGTC) and EukBr (5’**GTCTCGTGGGCTCGGAGATGTGTATAAGAGACAG**TGATCCTTCTGCAGGTTCACCTAC) from the Earth Microbiome Project (EMP) standard protocol (http://www.earthmicrobiome.org/protocols-and-standards/18s/), with Illumina Nextera adapters attached (adapters in bold). The mammalian blocking primer (5’GCCCGTCGCTACTACCGATTGG/ideoxyI//ideoxyI//ideoxyI//ideoxyI//ideoxyI/TTAGT GAGGCCCT/3SpC3/) was also included to reduce amplification of human 18S rRNA. Samples were amplified in duplicate with the PCR mixture and conditions specified in the EMP protocol, and duplicates were pooled after amplification. These amplicons were then sent to AGRF for purification, dual-indexing, pooling and sequencing.

### Sequence analysis

Demultiplexed sequence data was received from AGRF in FASTQ format and imported into the QIIME2 v2018.11 [24] version of the QIIME software [25] for analysis. Full documentation of all analysis code in this project can be found at https://rachaellappan.github.io/VL-QIIME2-analysis/. In brief, for both the 16S rRNA and 18S rRNA data, amplicon primers were first removed with the cutadapt plugin [26]. Reads were denoised, filtered and trimmed (where the median quality score fell below 30) with the DADA2 plugin [27]. Amplicon sequence variants (ASVs) were classified within QIIME2 using the SILVA v132 database [28], with a classifier trained on the amplified region [29]. These ASVs equate to classifying operational taxonomic units (OTUs) on the basis of 100% sequence identity. Note that where we use the traditional terminology of OTUs this differs from the traditional method of clustering sequences and calling OTUs based (usually) on 97% sequence identity [30]. The majority taxonomy with 7 levels was used for both the 16S rRNA and 18S rRNA datasets. *De novo* phylogenetic trees [31, 32] used in downstream measures of diversity that included phylogenetic distances (cf. below) were created using log_10_ read counts and the phylogeny align-to-tree-mafft-fastree (MAFFT multiple sequence alignment program) plugin in QIIME2. Stacked bar plots of percent relative abundance of different taxa across samples were generated in QIIME2. Aggregate percent abundance of taxa at the population level was determined by summing read counts per taxon across all study participants and calculating these as a proportion of the total read count. Prior to calculating downstream measures of diversity 18S rRNA data was also filtered to remove a small number of mammalian 18S rRNA reads, and also to remove bacterial 16S rRNA reads due to non-specificity of primers for eukaryotic 18S rRNA as described in the results section.

### Measures of diversity

Alpha (within-sample) diversity was measured using Shannon’s Diversity [33], Faith’s Phylogenetic Diversity (PD) [34], and Pilou’s Evenness [35] indices, calculated per sample within QIIME2 using rarefied counts (i.e. subsampled to the same sequencing depth across samples; rarefied to 21,383 for the 16S dataset, and 2,319 for the filtered 18S dataset). Boxplot figures for alpha diversity were created using GraphPad Prism v8, with between group statistical differences determined using the Mann-Whitney or ANOVA nonparametric tests in Prism. Beta (across-sample) diversity carried out using rarefied counts (as above) was measured using the Bray-Curtis dissimilarity statistic [36] based on compositional dissimilarity between samples taking abundance into account, unweighted UniFrac distances that measures phylogenetic distances between taxa, or weighted UniFrac distances [37] that measures phylogenetic distances and also takes relative abundance into account, as indicated. Principal coordinates (PCoA) plots for these indices were either generated within QIIME2 using the EMPeror graphics tools [38], or in QIIME1 version 1.9.1 [25] using the make_2d_plots.py script. Analysis of composition of microbiomes (ANCOM) [39], previously reported [40] as a sensitive method (for samples >20) with good control of false discovery rate, was also used within QIIME2 to determine between group differences in taxon abundance. The use of log ratios within this test takes account of differences in sequencing depth. Significance is reported as a W-value which provides a measure of how significant a specific taxon is relative to a number of other taxa used in the analysis.

## Results

### Prokaryotic and eukaryotic taxa identified across the study sample

A total of 12.5 million raw paired-end reads were generated across 46 faecal samples (mean±SD 271,509±166,138 reads/sample) for 16S rRNA amplicons, and a total of 1.6 million raw paired-end reads (mean±SD 34,112±6,852 reads/sample) for 18S rRNA amplicons. After read pre-processing of 16S rRNA sequence data there were 91,923±69,706 joined paired-end reads (minimum read depth 21,383; 33±6% of raw paired-end reads retained) per sample across VL and EC samples. Rarefaction plots (S1 Fig (a)) based on Shannon’s diversity index confirmed equivalent alpha diversity across the range of read depths from 2,000 to >20,000 (i.e. maximum alpha diversity is achieved at 2,000 reads). Using this read pre-processed data, 348 unique genera were identified from the 16S rRNA reads classified as ASVs within DADA2. The full taxonomy of these prokaryote 100% sequence identity OTUs is provided in S2 Table. For 18S rRNA the initial pre-processed read depth across experimental samples ranged from 21,460-49,822 joined paired-end reads (95±2% of raw paired-end reads retained). However, preliminary ASV classification demonstrated high abundance of unclassified “Eukaryota” across all samples (S2 Fig (a-b)). BLAST analysis of these unclassified taxa revealed alignment to bacterial 16S rRNA genes, demonstrating that the EMP primers for 18S rRNA amplification were not eukaryote-specific. These sequences were filtered out, leaving 77-27,315 joined paired-end reads per sample across the 46 samples. Using this filtered dataset, 62 unique genera were identified from the 18S rRNA reads classified as ASVs within DADA2. The full taxonomy of these eukaryotic 100% sequence identity OTUs is provided in S3 Table. Shannon’s index rarefaction plots (S1 Fig (b)) indicated equivalent alpha diversity across read depths ≥400 (i.e. maximum alpha diversity that it was possible to determine in our samples was achieved at ≥400 reads). However, to provide robustness in downstream analyses of diversity a rarefied read depth of 2,319 was used. This resulted in loss of N=8 VL and N=7 EC samples for whom read depths were <1000 reads, leaving sample sizes of 15VL and 16EC in the downstream eukaryotic analyses.

Fig 1(a) provides a bar plot of 16S rRNA-identified bacterial taxa ordered by abundance across the 46 samples. Fig 1(b) provides the bar plot for the mock positive controls (see also S2 Fig (c) for negative controls), demonstrating equivalent relative abundance of control bacterial taxa from the ZymoBIOMICS Microbial Community DNA Standard in the presence of increasing concentrations of human DNA. Table 2A shows the aggregated relative abundance at genus-level (% total counts and family provided in parentheses) for the 8 most prevalent taxa identified in the study sample. This includes *Prevotella* 9 (33.42%; Prevotellaceae), *Faecalibacterium* (11.32%; Ruminococcaceae), *Escherichia-Shigella* (9.14%; Enterobacteriaceae), *Alloprevotella* (4.46%; Prevotellaceae), *Bacteroides* (4.07%; Bacteroidaceae), *Prevotella* 2 (3.61%; Prevotellaceae), *Ruminococcaceae* UCG-002 (1.58%; Ruminococcaceae), and *Bifidobacterium* (1.54%; Bifidobacteriaceae). Fig 1(c) provides a heatmap of relative abundances for the top genera (*Prevotella*, *Faecalibacterium*, *Bacteroides* and *Escherichia-Shigella*) that relate to previously described enterotypes discussed in more detail in the discussion section below [16, 41]. Our study sample suggests a predominance of enterotype-2 driven by *Prevotella*, with only a small number of individuals conforming to enterotype-1 driven by *Bacteroides* and a lack of enterotype-3 driven by *Ruminococcus*. High *Faecalibacterium* abundance was generally associated with high *Prevotella*, whereas *Escherichia-Shigella* was associated with low *Prevotella*. Bacterial taxa (e.g. *Dialister*, *Megasphaera*, *Mitsuokella*, *Lactobacillus*) previously shown to be characteristic of Indian gut microbiomes [17, 21, 42] occurred at <1% in our study population (Table 2).

**Fig 1.**
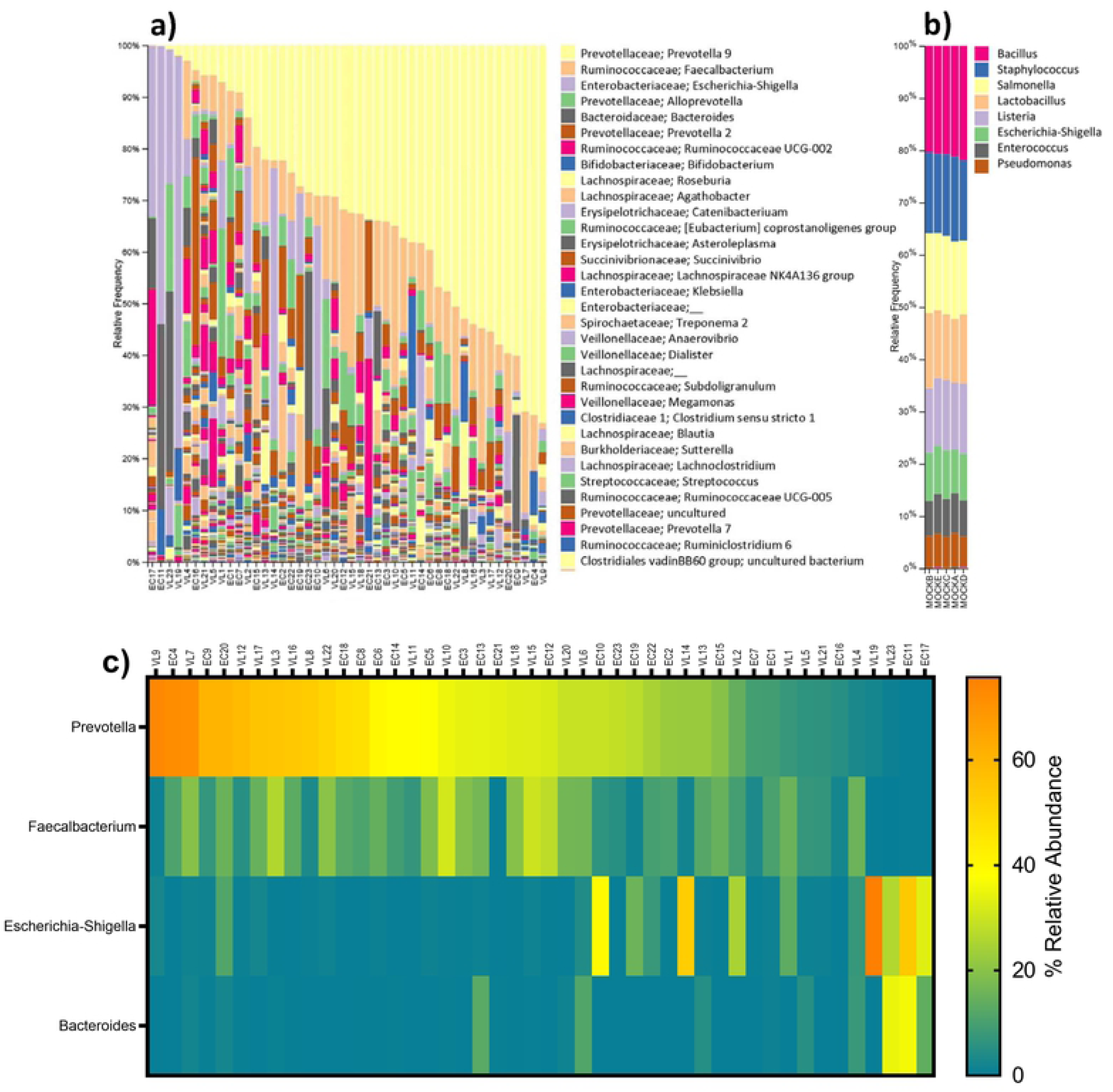
Results of 16S rRNA analysis of ASVs at genus level in the study sample. (a) Bar plot ordered by relative abundance for prokaryotic taxa colour coded according to the key to the right of the bar plot. (b) Bar plot for prokaryotic mock control samples. (c) Heat map of relative abundance for the 4 taxa associated with previously defined bacterial enterotypes, ordered by relative abundance of *Prevotella*.

**Table 2.** Genus-level community composition, and family to which they belong, for VL cases, endemic controls, and the total study sample. Abundance at the study group level was calculated for aggregated read counts across all samples for ASVs classified at genus level (i.e. aggregates all taxa with the same genus assignment). Genera with an average group-level abundance <1% are not shown, except where relevant to the results or discussion. Full details are provided in S2 and S3 Tables.

| Genus level taxonomy | Family level taxonomy | Percent in VL cases | Percent in endemic controls | Percent across all samples |
| --- | --- | --- | --- | --- |
| **A. Prokaryotes based on 16S rRNA sequencing** |  |  |  |  |
| *Prevotella* (2+7+9) | Prevotellaceae | 40.77 | 33.61 | 37.47 |
| *Faecalibacterium* | Ruminococcaceae | 12.44 | 10.02 | 11.32 |
| *Escherichia-Shigella* | Enterobacteriaceae | 8.82 | 9.51 | 9.14 |
| *Alloprevotella* | Prevotellaceae | 4.31 | 4.63 | 4.46 |
| *Bacteroides* | Bacteroidaceae | 3.46 | 4.78 | 4.07 |
| *Ruminococcaceae* UCG-002 | Ruminococcaceae | 1.87 | 1.24 | 1.58 |
| *Bifidobacterium* | Bifidobacteriaceae | 1.67 | 1.38 | 1.54 |
| *Roseburia* | Lachnospiraceae | 0.68 | 2.38 | 1.47 |
| *Agathobacter* | Lachnospiraceae | 0.47 | 2.43 | 1.38 |
| *Catenibacterium* | Erysipelotrichaceae | 0.73 | 1.76 | 1.21 |
| *Asteroleplasma* | Erysipelotrichaceae | 0.02 | 2.45 | 1.14 |
| *Succinivibrio* | Succinivibrionaceae | 0.71 | 1.38 | 1.02 |
| *Clostridiales vadin BB60 group* | Order: Clostridiales | 1.38 | 0.31 | 0.89 |
| *Anaerovibrio* | Veillonellaceae | 0.01 | 1.48 | 0.69 |
| *Dialister* | Veillonellaceae | 0.41 | 0.91 | 0.64 |
| *Megasmonas* | Veillonellaceae | 0.25 | 0.99 | 0.59 |
| *Megasphaera* | Veillonellaceae | 0.57 | 0.14 | 0.37 |
| *Mitsuokella* | Veillonellaceae | 0.07 | 0.13 | 0.10 |
| *Lactobacillus* | Lactobacillaceae | 0.10 | 0.08 | 0.09 |
| *Ruminococcaceae* UCG-014 | Ruminococcaceae | 0.05 | 0.28 | 0.15 |
| *Gastranaerophilales uncultured bacterium* | Order: Gastranaerophilales | 0.002 | 0.24 | 0.15 |
| **B. Eukaryotes based on 18S rRNA sequencing** |  |  |  |  |
| *Blastocystis* sp. MJ99-568 | Order: Stramenopiles | 38.88 | 25.48 | 32.27 |
| *Blastocystis* ambiguous taxon | Order: Stramenopiles | 21.32 | 29.78 | 25.49 |
| *Dientamoeba* | Trichomonadea | 9.38 | 14.94 | 12.12 |
| *Pentatrichomonas* | Trichomonodea | 18.76 | 1.29 | 10.13 |
| *Entamoeba* | Entamoebida | 0.92 | 6.04 | 3.45 |
| Ascaridida | Chromodorea | 0 | 1.58 | 0.78 |
| Rhabditida | Chromodorea | 0.86 | 0.69 | 0.77 |
| Cyclophyllidea | Eucestoda | 0 | 0.38 | 0.19 |

Fig 2(a) provides a bar plot of 18S rRNA-identified eukaryotic taxa ordered by abundance across the 46 samples. Fig 2(b) provides the bar plot for the mock positive controls, demonstrating equivalent relative abundance of the two control fungal eukaryotes in the ZymoBIOMICS Microbial Community DNA Standard, together with identification of *Leishmania* and *Toxoplasma* DNA spiked into the control samples in-house. Full details of positive and negative control samples are provided in S2 Fig. Table 2B shows the aggregated relative abundance at genus or lowest order level classified (higher order classification also provided in parentheses) for the eukaryotic taxa of interest (i.e. excluding plant genera) identified in the study sample. This includes *Blastocystis* sp. MJ99-568 (32.37%; Order: Stramenopiles), *Blastocystis* ambiguous taxon (25.49%; Order: Stramenopiles), *Dientamoeba* (12.12%; Family: Tritrichomonadea), *Pentatrichomonas* (10.13%; Family: Trichomonodea), *Entamoeba* (3.45%; Family: Entamoebida), Ascaridida (0.78%; Family: Chromodorea; likely *Ascaris* round worm cf. below), Rhabditida (0.77%; Family: Chromodorea; likely *Strongyloides* or hookworm cf. below), and Cyclophyllidea (0.19%; Order: Eucestoda; likely *Hymenolepis* tapeworm cf. below). Of these, unsupervised cluster analysis (Fig 2(c)) shows 3 clusters of individuals that are defined by high relative frequencies of: (A) N=10 (3 VL; 7 EC) *Blastocystis* Ambiguous taxon with/without *Dientamoeba* and/or *Entamoeba*; (B) N=11 (9 VL; 2 EC) with *Pentatrichomonas* and/or *Blastocystis* sp. MJ99-568 in the absence of taxa from cluster A; and (C) N=10 (3VL; 7EC) absence or low abundance of all 5 of these taxa but high frequency of *Saccharomcyes* and plant species. We refer to these below as “eukaryotic enterotypes” A, B and C. Species level identification in QIIME2 confirmed *Pentatrichomonas* as *P. hominis* and *Dientamoeba* as *D. fragilis*. *Entamoeba* was not identifiable to species level.

**Fig 2.**
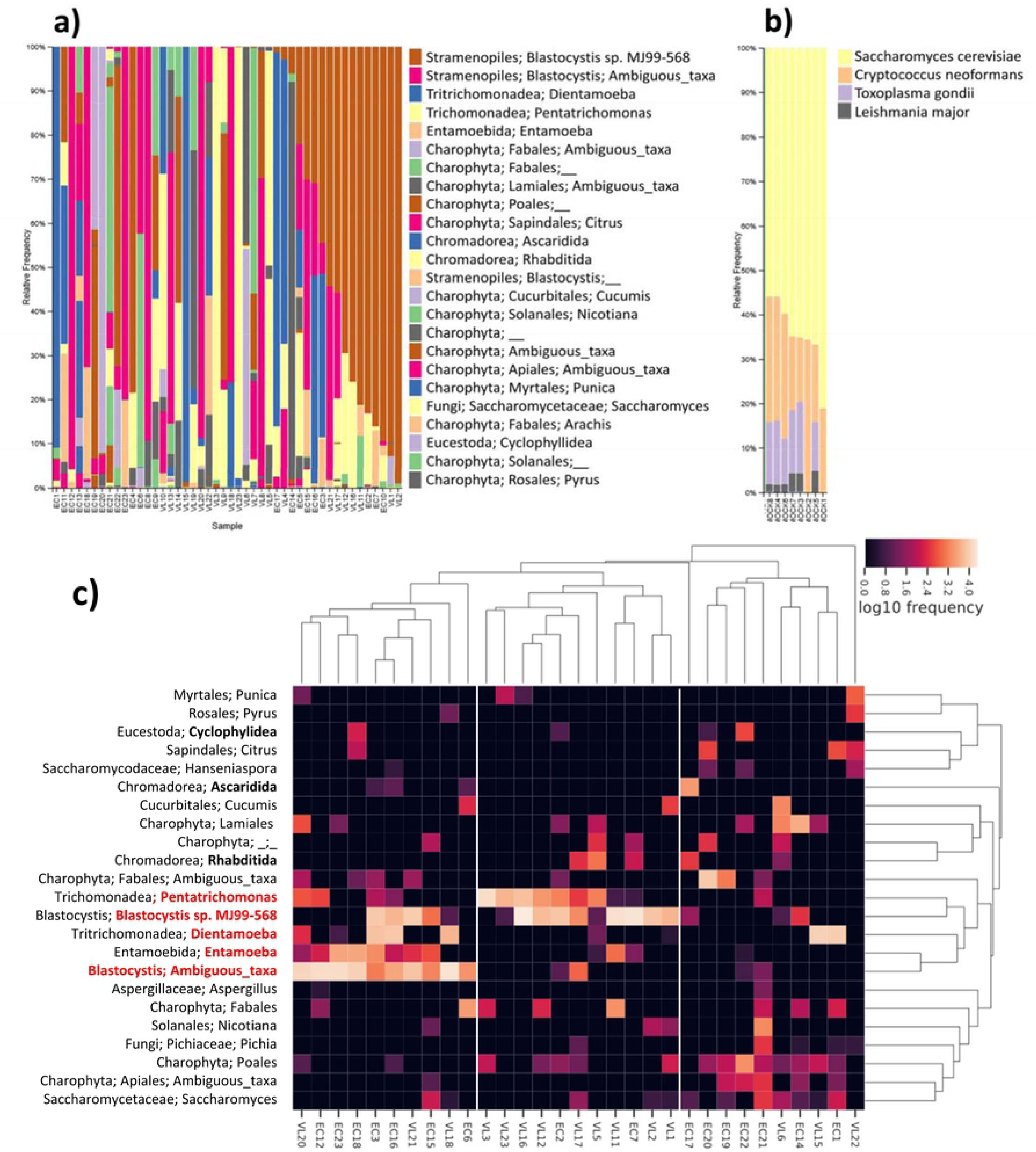
Results of 18S rRNA analysis of ASVs at genus level in the study sample. (a) Bar plot ordered by relative abundance for eukaryotic taxa colour coded according to the key to the right of the bar plot. (b) Shows the bar plot for eukaryotic mock control samples. (c) Heatmap derived from unsupervised cluster analysis of eukaryotic taxa. Red labelling indicates the cluster of 5 most prevalent eukaryotic protozoan taxa for which samples cluster into three eukaryotic enterotypes according to: (A) *Blastocystis* Ambiguous taxon with/without *Dientamoeba* and/or *Entamoeba*; (B) *Pentatrichomonas* and/or *Blastocystis* sp. MJ99-568 in the absence of taxa from cluster A; (C) low abundance of these protozoan taxa. Bold labelling indicates the helminth eukaryotic taxa. Other eukaryotic taxa include fungi and a variety of plant taxa derived from the diet of participants.

### Influence of age, sex or microgeographic location on gut microflora in the study sample

Other studies in India [18, 20] and elsewhere [15, 41] have demonstrated changes in the composition of gut flora with age. In our study it was important to determine whether there were any global effects of age and/or gender on microbial diversity that needed to be adjusted for in analysis of differences between VL cases and EC. S3 Fig (a-b) provide bar plots of the 16S rRNA data reordered by age or gender (and secondarily on relative abundances of taxa; key as per Fig 1(a)), respectively. Visual inspection suggests no overt effects of either age or sex on composition of microbial flora. This is confirmed comparing species richness (Number of ASVs ≅ 100% OTUs) and alpha diversity measures (Faith’s PD, Pielou Evenness, Shannon’s diversity index) between genders (S3 Fig (c-f)) and beta diversity PCoA plots for weighted UniFrac distances by gender (S3 Fig (g)) and for age as a continuous variable (S3 Fig (h)). Similarly, S4 fig (a-b) provide bar plots of the 18S rRNA data reordered by age and gender, respectively. Again, visual inspection suggests no overt effects of either age or sex on composition of microbial flora. This is supported by comparisons of species richness (Number of ASVs ≅ 100% OTUs) and alpha diversity measures (Faith’s PD, Pielou Evenness, Shannon’s diversity index) between sexes (S4 Fig (c-f)) and PCoA plots for weighted UniFrac distances by gender (S4 Fig (g)) and for age as a continuous variable (S4 Fig (h)). In this case there was borderline significance (p=0.048) for difference between genders for Faith’s phylogenetic diversity. On balance, we conclude there are no overt independent effects of age or sex on either 16S rRNA-determined bacterial diversity or 18S rRNA-determined eukaryotic diversity in gut flora that would affect analysis of microbial profiles comparing VL cases with EC.

Substructure in microbial diversity due to microgeographic location could also confound downstream analysis comparing VL cases and EC. Generally, EC were matched to VL for location as they were attendants of patients at KAMRC. KAMRC is a centre that draws patients from across Bihar State in India, but principally (in this study) from within the district of Muzzaffapur. S5 Fig (a) provides a map of subdistricts or blocks within Muzzaffapur District showing participant locations. S5 Fig (b) shows the 16S rRNA relative abundance bar plot reordered by subdistrict (key to taxa as per Fig 1(a)). Box plots (S5 Fig (c-f)) comparing species richness (Number of ASVs ≅ 100% OTUs) and alpha diversity measures (Faith’s PD, Pielou Evenness, Shannon’s diversity index) across blocks where N≥4 indicate no statistically significant differences by subdistrict. Principle coordinate plots based on weighted UniFrac distances (S5 Fig 5(g)) also show no overt clustering by subdistrict. Similar results were obtained for the 18S rRNA data (S6 Fig). We conclude there are no overt differences in gut microflora by subdistrict in this region of Bihar State.

### Comparing gut microflora between VL cases and EC in the study sample

Fig 3(a-b) provide bar plots of the relative abundances of taxa reordered by VL case or EC status for 16S rRNA data and 18S rRNA data, respectively. Species richness (Number of ASVs ≅ 100% OTUs) and alpha diversity box plots (Faith’s PD, Pielou Evenness, Shannon’s diversity index) comparing VL cases with EC (Fig 3(c-f) for 16S rRNA; Fig 3(g-j) for 18S rRNA) show no statistically significant differences in alpha diversity between these two groups. Beta-diversity PCoA plots based on Bray-Curtis distances (Fig 4(a-b)) that take relative abundance into account or using weighted UniFrac distances (Fig 4(c-d)) that take abundance and phylogenetic distances between taxa into count, also show no clear evidence for clustering based on VL cases *versus* EC status.

**Fig 3.**
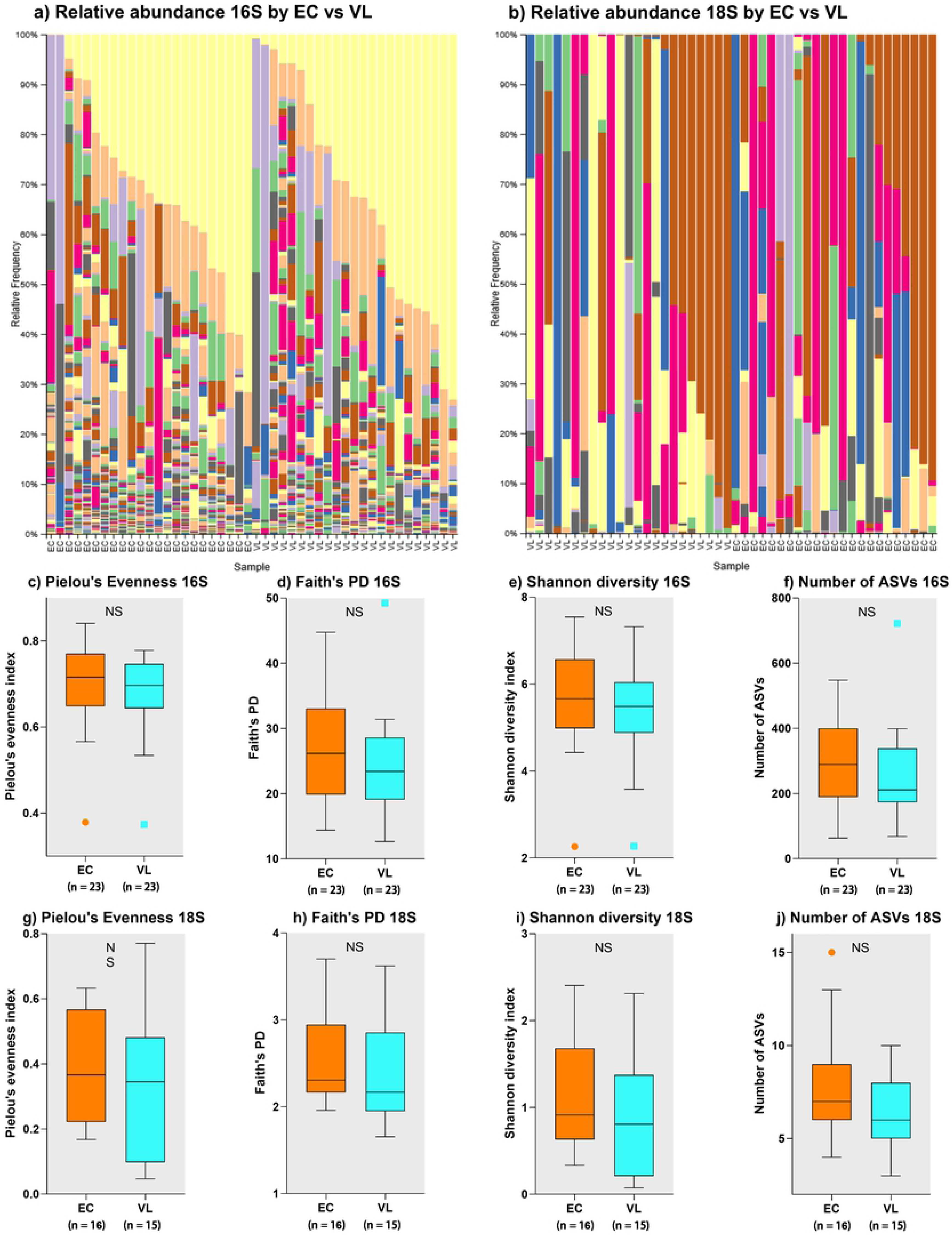
Analysis of gut microbial profiles in relation to VL case versus EC status. on 16S rRNA-determined prokaryotic and 18S rRNA-determined eukaryotic gut microbial profiles. (a) bar plots for relative abundance of 16S rRNA taxa separated for VL cases and EC groups, each ordered by relative abundance of taxa; for colour coding see key to main Fig 1(a). (b) bar plots for relative abundance of 18S rRNA taxa separated for VL cases and EC groups, each ordered by relative abundance of taxa; for colour coding see key to main Fig 2(a). (c) to (f) comparisons of species richness (number of ASVs) and alpha diversity measures (as labelled) for 16S rRNA data. (g) to (j) comparisons of species richness (number of ASVs) and alpha diversity measures (as labelled) for 18S rRNA data.

**Fig 4.**
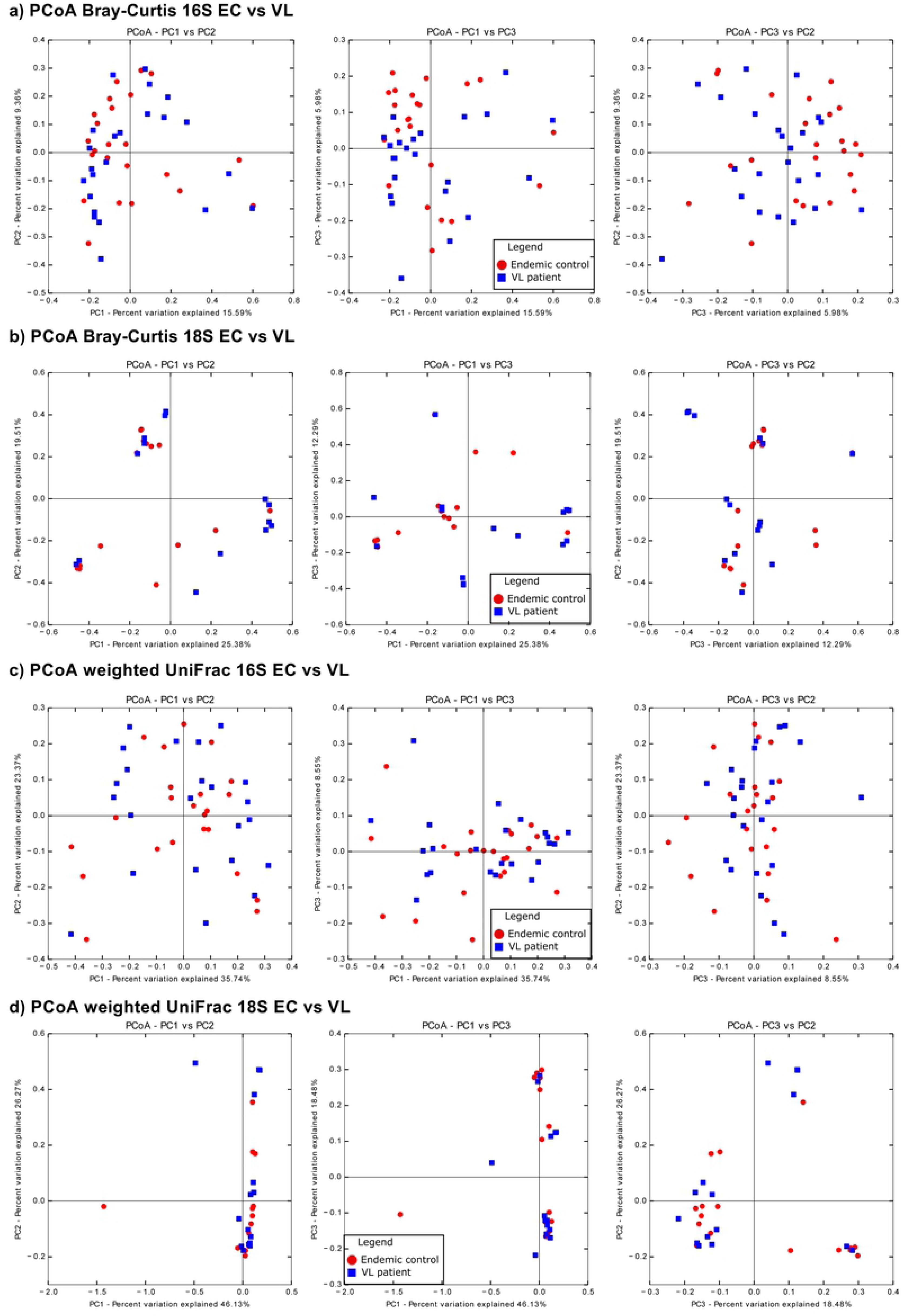
Beta diversity comparisons for VL cases and EC. (a) and (b) are PCoA plots (left to right: PC1xPC2; PC1xPC3; PC2xPC3) for Bray-Curtis dissimilarity measures for 16S rRNA and 18S rRNA data, respectively. (c) and (d) are PCoA plots (left to right: PC1xPC2; PC1xPC3; PC2xPC3) for weighted UniFrac measures for 16S rRNA and 18S rRNA data, respectively. See keys on plots for colour coding of VL cases and EC. Note: the weighted UniFrac distances are skewed due to 2 samples with high worm burdens.

Although there were no global differences in gut prokaryotic or eukaryotic microbial composition between VL cases and EC as determined by alpha and beta diversity measures, analysis of 16S rRNA data using ANCOM identified two bacterial genera, Ruminococcaceae UCG-014 (Family: Ruminococcaceae) and Gastranaerophilales_uncultured bacterium (Phylum: Cyanobacteria; Class: Melainabacteria; Order Gastranaerophilales), that were enriched in EC compared to VL cases (Fig 5 (a-b)). These differences are also apparent in the aggregated relative abundance data presented in Table 2, with Ruminococcaceae UCG-014 at 0.28% in EC compared to 0.05% in VL cases and Gastranaeophilales_uncultured at 0.24% in EC compared to 0.002% in VL cases.

**Fig 5.**
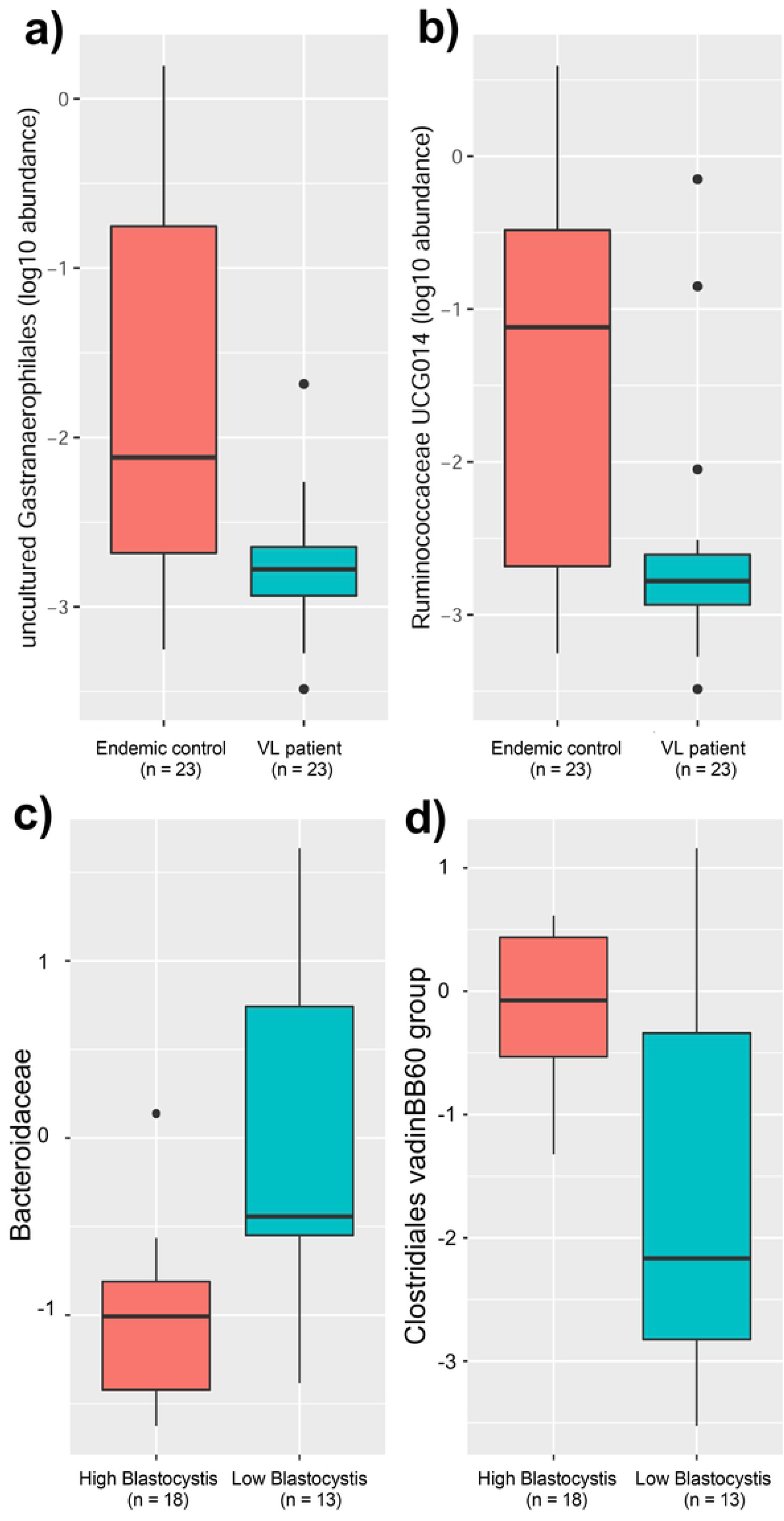
Results of ANCOM analysis of between-group differences in microbial composition. (a) and (b) box plots for prokaryotic taxa that differed between EC and VL cases. W-values were 41 for Gastranaerophilales presented in (a) and 61 for Ruminococcaceae presented in (b), indicating that these taxa were significantly higher in the EC group compared to VL in comparison with 41 or 61 other taxa examined, respectively. (c) and (d) ANCOM box plots for prokaryotic taxa that differed between groups of individuals carrying high (46-89%) versus low (<6%) relative abundance of eukaryotic Blastocystis. W-value = 31 (c) indicating that Bacteroides abundance was significantly lower in the high Blastocystis group in comparison with 31 other bacterial families examined. W-value = 30 (d) indicating that *Clostridiales vadinBB60* abundance was significantly higher in the high Blastocystis group in comparison with 30 other bacterial families examined.

Visual inspection of the 18S rRNA bar plots (Fig 2(a)) suggested a trend for higher abundance of *P. hominis* in VL cases compared to EC. In contrast, there were more individuals with higher relative abundance of *Entamoeba* in EC than in VL cases. This is also reflected in the aggregated relative abundance data for eukaryotic genera presented in Table 2B, and further highlighted on a heatmap (Fig 6(a)) for eukaryotic protozoa log_10_ read counts ordered by *P. hominis* counts in EC and VL cases. This aligns with the “eukaryotic enterotypes” defined above, where 7/9 individuals classified as “eukaryotic enterotype B” defined by relative frequencies of read counts for *P. hominis* and *Blastocystis* sp. MJ99-568 were VL patients, and conversely 7/10 individuals classified as “eukaryotic enterotype A” defined by *Blastocystis* ambiguous taxon, with/without *Entamoeba* or *D. fragilis*, were EC. Comparisons of log_10_ read counts (Fig 6(b-c)) and relative abundance (Fig 6(d-e)) show that VL cases positive for *P. hominis* had significantly higher parasite burdens compared to EC, whereas this was not the case for the between group comparison of individuals positive for *Entamoeba*. This suggests that when VL patients acquire *P. hominis* they have a more intense colonisation whereas when they acquire *Entamoeba* the relative abundance is the same as for EC individuals.

**Fig 6.**
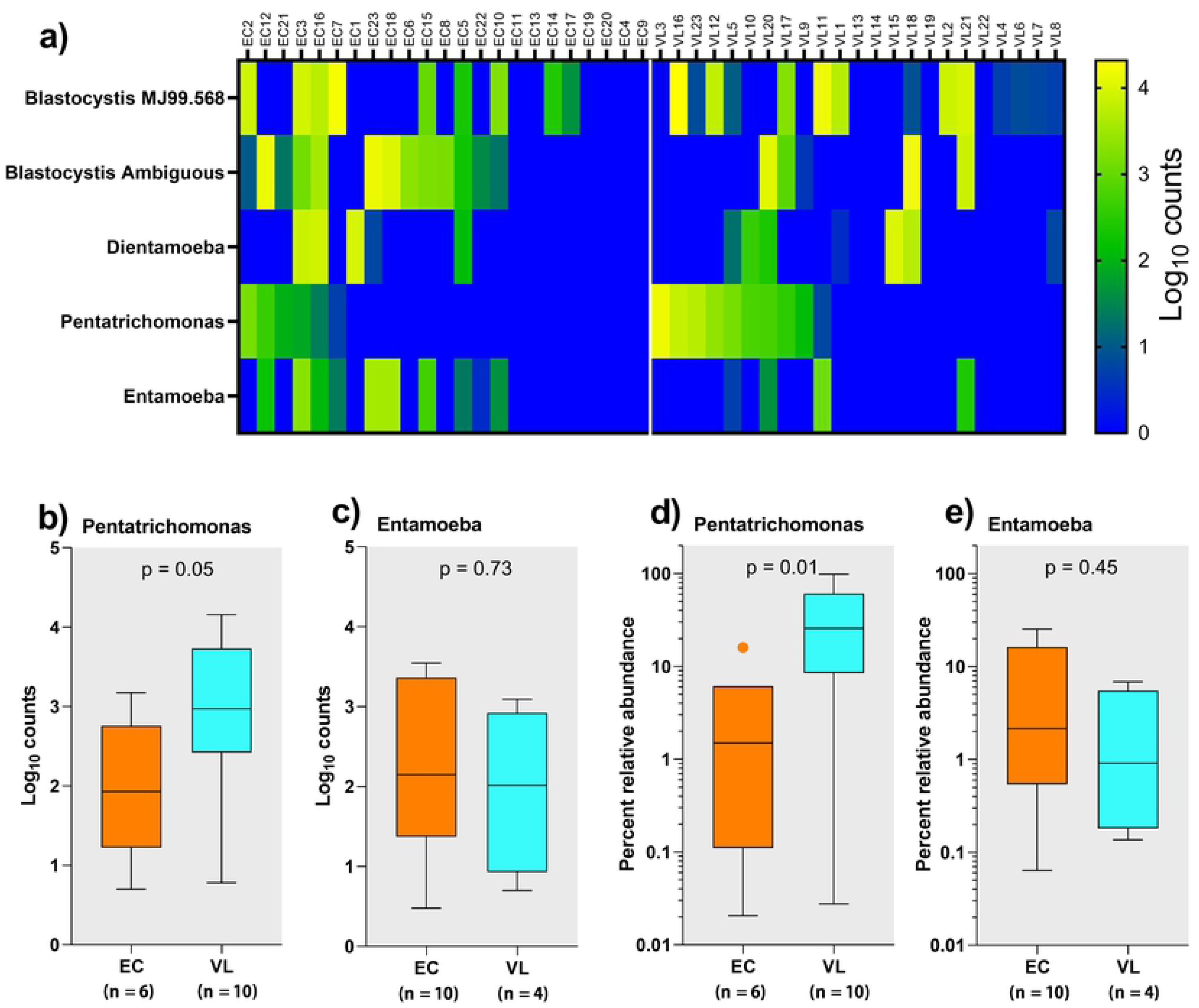
Influence of VL on eukaryotic *Pentatrichomonas* burden. (a) heatmap of the top 5 eukaryotic taxa separated by VL case versus EC status, each ordered according to *Pentatrichomonas* load. (b) and (c) box plots comparing *Pentrichomona*s and *Entamoeba* log_10_ counts between EC and VL cases. (d) and (e) show the same comparisons for percent relative abundance. VL cases carry higher loads of *Pentatrichomonas* but not of *Entamoeba*.

### Influence of known enteric pathogens on microbial diversity in gut flora

Entero-pathogens have been observed to perturb bacterial diversity in gut flora, for example in the context of a bacterial enterotype defined by *Escherichia-Shigella* abundance [41]. We therefore looked across our sample to see if such effects could be observed in our study population. Looking across relative abundance plots for 16S rRNA data (see Fig 1 and Fig 3) for VL cases and EC there were no overt group-specific differences in relative abundance of bacterial taxa that could contain pathogenic species, e.g. *Escherichia-Shigella*. However, some individuals positive for *Escherichia-Shigella* did appear to have reduced diversity in terms of other bacterial species in their microflora. To quantify this, we compared alpha (Pielou’s Evenness, Faith’s PD, Shannon’s Diversity) diversity measures for individuals who were negative or had very low (<7.5%; N=33) relative abundance of *Escherichia-Shigella* with those who had high relative abundance (>11.3%; N=10) (Fig 7(a-c)). Significant differences in Pielou’s evenness (p=0.0007) and Shannon’s diversity (p=0.002), both of which take abundance and evenness of taxa into account, were observed. Faith’s PD, which takes phylogenetic distances into account, did not differ significantly between groups with high and low *Escherichia-Shigella* relative abundance. To ensure any effects observed weren’t due to artefact in using relative abundance data (i.e. if you have a lot of one taxon then there is bound to be reduced abundance of other taxa) we carried out a similar analysis of individual with low (<10%; N=11) or high (>14%; N=35) abundance of the dominant commensal genus *Prevotella 9* (Fig 7(d-f)). No significant differences in Pielou’s evenness index or Shannon’s diversity were observed, but there was borderline significance (p=0.04) for Faith’s PD between these two groups. Overall, we conclude that high abundance of *Escherichia-Shigella* is influencing dysbiosis of gut bacterial microflora.

**Fig 7.**
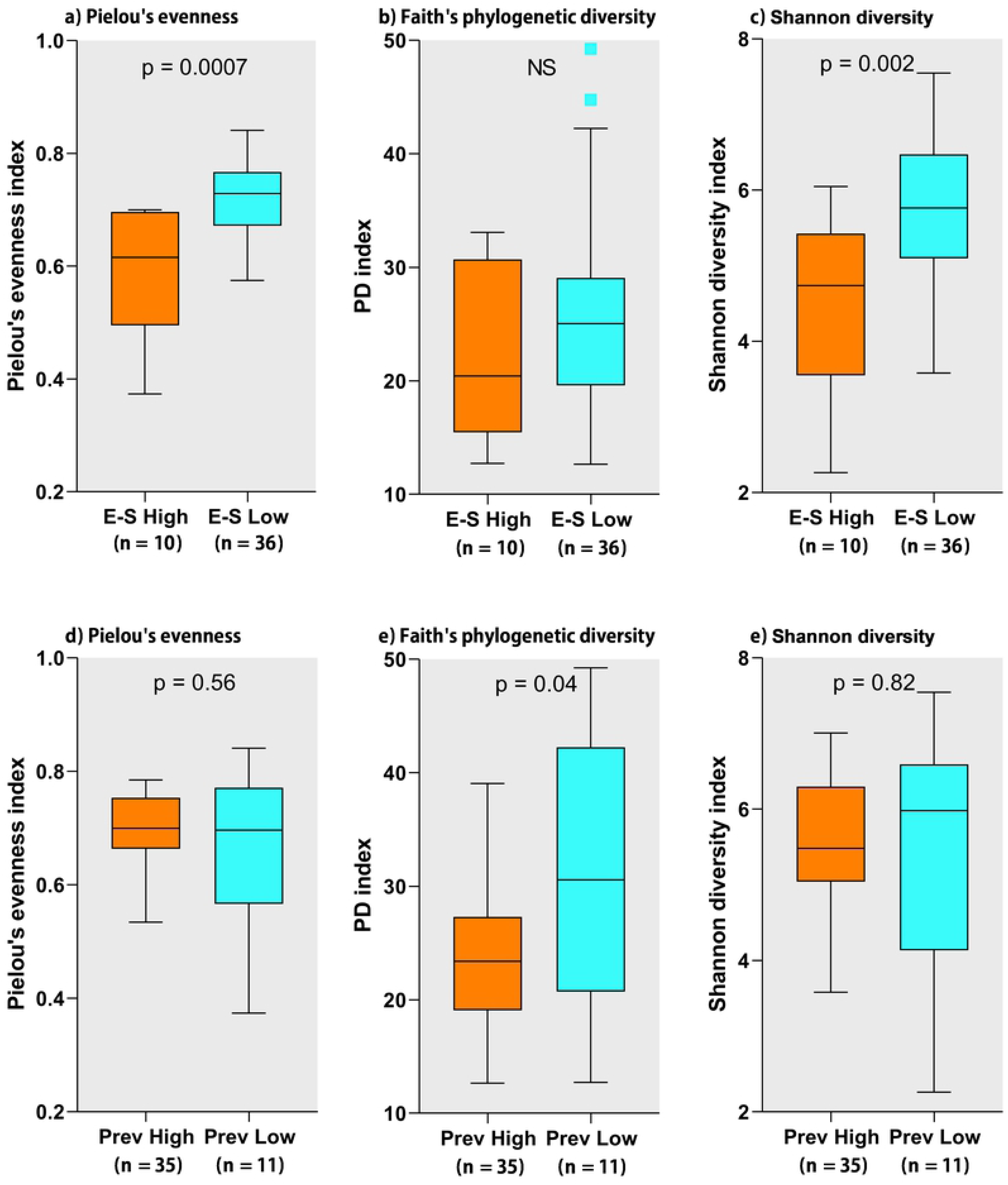
Influence of *Escherichia-Shigella* on alpha diversity of bacterial taxa. (a-c) box plots comparing 16S rRNA alpha diversity measures (as labelled) between groups of individuals carrying high or low *Escherichia-Shigella* loads. (d-f) box plots making the same comparisons for groups of individuals carrying high or low *Prevotella* 9 loads.

Another eukaryotic genus that contains potentially pathogenic species is *Blastocystis*. In looking for effects on the bacterial microflora of eukaryotic taxa that might contain putative pathogens, we noted clustering according to *Blastocystis* abundance on Jaccard index beta diversity PCoA plots (Fig 8(a-c)) and unweighted Unifrac PCoA plots (Fig 8 (d)). This was true for both *Blastocystis* sp. MJ99-568 (Fig 8(a)) and *Blastocystis*_ambiguous taxon (Fig 8(b)). Specifically, individuals with high *Blastocystis* abundance (46-98%) were almost exclusively clustered to the right of the zero co-ordinate on PCoA axis 1, whereas individuals with low Blastocystis abundance (<6%) were clustered to the left. To determine what bacterial taxa might be driving this variation we carried out biplot analyses, results of which are superimposed onto PCoA plots of Jaccard indices based on presence/absence of bacterial taxa (Fig 8(a-c)) and onto PCoA plots of unweighted UniFrac distances that take phylogenetic distances into account (Fig 8(d)). The four most significant bacterial taxa driven by eukaryotic *Blastocystis* abundance were *Escherichia-Shigella, Prevotella* 9, *Succinivibrio*, and *Asteroleplasma* for the Jaccard index analysis, and *Escherichia-Shigella, Bacteroides thetaiotaomicron, Faecalibacterium, Prevotella* 9 based on unweighted UniFrac distances. In both cases *Escherichia-Shigella* had the strongest effect (as indicated by the length of arrow on the biplots), which manifests as a negative correlation between log_10_ counts for *Escherichia-Shigella* and *Blastocystis* (Fig 8(e)). *Blastocystis* relative abundance also had a more general influence on alpha-diversity measures (Fig 9(a-c) for *Blastocystis* MJ99-568; Fig 9(e-g) for *Blastocystis* Ambiguous taxon) and on species richness (Number of ASVs ≅ 100% OTUs) (Fig 9(d) for *Blastocystis* MJ99-568; Fig 9(h) for *Blastocystis* Ambiguous taxon). Notably, high abundance of both *Blastocystis* taxa was associated with higher species richness and higher measures of alpha diversity. To further identify specific bacterial taxa that might be related to the effects of *Blastocystis* on bacterial diversity we employed ANCOM to look at the family-level for differential 16S rRNA-determined bacterial abundance between high versus low *Blastocystis* carriers. This identified two bacterial taxa associated with high versus low *Blastocystis* count (Fig 5(c-d)). Bacteroidaceae (aggregate abundance 4.07%, see Table 2) were less abundant with high *Blastocystis* and more abundant with low *Blastocystis* (Fig 5(c)). Conversely, *Clostridiales* vadinBB60 group (aggregate abundance 0.89%, see Table 2) were more abundant with high *Blastocystis* and less abundant with low *Blastocystis* (Fig 5(d)).

**Fig 8.**
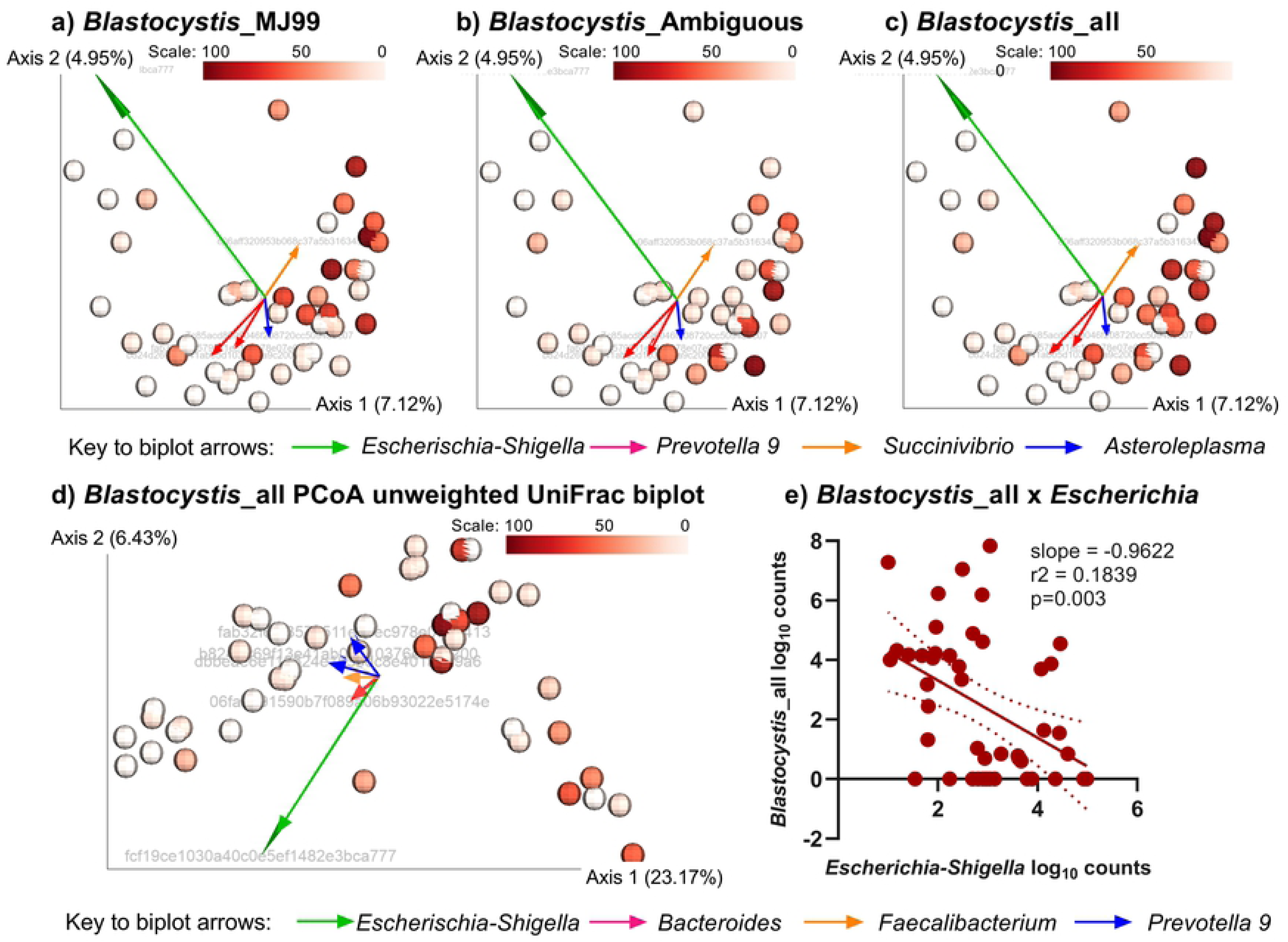
Influence of *Blastocystis* on beta diversity of bacterial taxa. (a-c) PCoA plots of Jaccard indices for beta diversity of bacterial microflora colour coded (see scales) according to relative abundances for (a) *Blastocystis*_MJ99-340, (b) *Blastocystis*_Ambiguous taxon, and (c) all *Blastocystis*. Results of the biplot analysis are superimposed on the PCoA plots, with length of arrows indicating the degree of influence of different bacterial taxa (see key to biplot arrows) on *Blastocysti*s abundance. (d) PCoA plot of unweighted UniFrac distances colour coded (see scale) according to *Blastocystis* load, with biplot vectors superimposed as before. (e) graph showing negative correlation between *Blastocystis* log_10_ counts and *Escherichia-Shigella* log_10_ counts.

**Fig 9.**
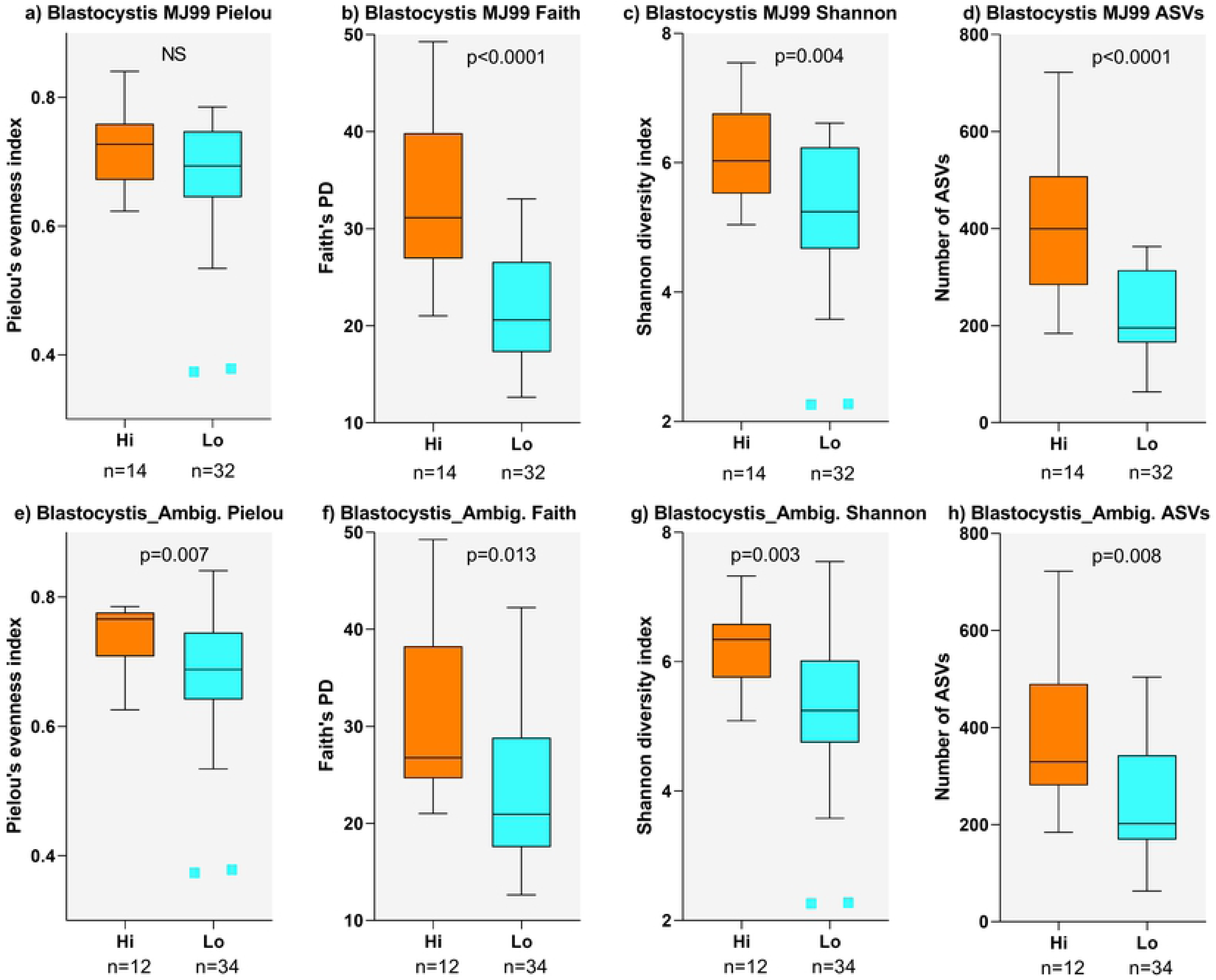
Influence of *Blastocystis* on alpha diversity of bacterial taxa. (a-d) box plots comparing influence of high or low *Blastocystis* MJ99-340 abundance on alpha diversity measures (as labelled) and species richness (as measured by number of ASVs). (e-f) the same comparisons for groups of individuals with high or low *Blastocystis*_Ambiguous taxon abundance.

### Using 18S rRNA analysis to identify gut helminths

One important reason for carrying out analysis of taxa based on 18S rRNA sequencing was to determine if it would be sufficiently sensitive to detect helminth species in faecal samples. Notwithstanding loss of read depth and sample numbers due to bacterial contamination of reads, we were able to detect 4 taxa containing pathogenic helminths, namely Ascaridida (*Ascaris*), Trichocephalida (*Trichuris*), Rhabditida (Hookworm – *Ancylostoma*) and Cyclophylidea (*Hymenolepis*). Table 3 compares data based on microscopy and 18S rRNA analysis for those individuals (N=36) for whom both datasets were available. Analysis using 18S rRNA identified 3/3 positive for *Ascaris* by microscopy, 0/8 positive for *Trichuris* by microscopy (with one that was positive for 18S rRNA but negative for microscopy), 7/7 positive for hookworm by microscopy (plus one positive by 18S rRNA and negative by microscopy), and 4/4 positive for *Hymenolepis* by microscopy.

**Table 3.** Comparison of helminth identification by microscopy (yes/no for presence/absence) with log_10_ read counts for parallel identification to the appropriate order using 18S rRNA analysis. Bold indicates where positive microscopy and 18S rRNA concur; italics indicates where they do not concur.

| Sample number | Ascaris | Ascaridida | Trichuris trichura | Trichocephalida | Hookworm | Rhabditida | Hymenolepis | Cyclophyllidea |
| --- | --- | --- | --- | --- | --- | --- | --- | --- |
| EC01 | No | 0 | No | 0 | No | 0 | No | 0 |
| EC02 | No | 0 | No | 0 | No | 0 | **Yes** | **0.90** |
| EC03 | **Yes** | **0.78** | No | 0 | No | 0 | No | 0 |
| EC04 | No | 0 | No | 0 | **Yes** | **2.44** | No | 0 |
| EC05 | No | 0 | *Yes* | *0* | **Yes** | **2.43** | No | 0 |
| EC06 | **Yes** | **1** | *Yes* | *0* | No | 0 | No | 0 |
| EC08 | No | 0 | *Yes* | *0* | No | 0 | No | 0 |
| EC09 | No | 0 | No | 0 | No | 0 | No | 0 |
| EC10 | No | 0 | No | 0 | **Yes** | **2.06** | No | 0 |
| EC11 | No | 0 | No | 0 | No | 0 | No | 0 |
| EC12 | No | 0 | No | 0 | No | 0 | No | 0 |
| EC14 | No | 0 | No | 0 | No | 0 | No | 0 |
| EC15 | No | 0 | No | 0 | No | 0 | No | 0 |
| EC17 | **Yes** | **3.41** | No | 0 | **Yes** | **2.52** | No | 0 |
| EC18 | No | 0 | No | 0 | No | 0 | **Yes** | **2.24** |
| EC19 | No | 0 | No | 0 | No | 0 | No | 0 |
| EC20 | No | 0 | No | 0 | No | 0 | **Yes** | **0.78** |
| EC21 | No | 0 | No | 0 | No | 0 | No | 0 |
| EC22 | No | 0 | No | 0 | No | 0 | **Yes** | **2.64** |
| EC23 | No | 0 | No | 0 | No | 0 | No | 0 |
| VL01 | No | 0 | No | 0 | No | 0 | No | 0 |
| VL02 | No | 0 | No | 0 | No | 0 | No | 0 |
| VL03 | No | 0 | No | 0 | No | 0 | No | 0 |
| VL04 | No | 0 | No | 0 | No | 0 | No | 0 |
| VL05 | No | 0 | *Yes* | *0* | **Yes** | **3.04** | No | 0 |
| VL06 | No | 0 | No | 0 | **Yes** | **1.41** | No | 0 |
| VL08 | No | 0 | No | 0 | No | 0 | No | 0 |
| VL09 | No | 0 | *Yes* | *0* | No | 0 | No | 0 |
| VL11 | No | 0 | *Yes* | *0* | No | 0 | No | 0 |
| VL12 | No | 0 | *No* | *0.60* | No | 0 | No | 0 |
| VL14 | No | 0 | No | 0 | **Yes** | **1.34** | No | 0 |
| VL16 | No | 0 | *Yes* | *0* | No | 0 | No | 0 |
| VL17 | No | 0 | *Yes* | *0* | *No* | *2.42* | No | 0 |
| VL20 | No | 0 | No | 0 | No | 0 | No | 0 |
| VL21 | No | 0 | No | 0 | No | 0 | No | 0 |
| VL22 | No | 0 | No | 0 | No | 0 | No | 0 |

## Discussion

In this study we have used a meta-taxonomic approach to determine the composition of prokaryotic and eukaryotic gut microflora in a region of Bihar State in North-East India endemic for VL. Amplicon-based sequencing of the V3-V4 region of the 16S rRNA gene showed that the most abundant bacterial taxa identified in faecal samples from this region of India were *Prevotella* (37.1%), *Faecalibacterium* (11.3%), *Escherichia-Shigella* (9.1%), *Alloprevotella* (4.5%), *Bacteroides* (4.1%), *Ruminococcaceae* UCG-002 (1.6%), and *Bifidobacterium* (1.5%). Our study was also unique in using alternative primers for amplicon-based sequencing of the V9 region of the 18srRNA gene to identify eukaryotic microflora. Although we encountered one issue with cross-amplification of 16S rRNA sequences due to sequence similarity of primers, we were successful in (a) blocking amplification of human 18S rRNA, and (b) in amplifying protozoan and metazoan sequences of pathogenic interest. In so doing we determined that the most prevalent protozoan taxa present in this region of India were *Blastocystis* sp. MJ99-568 (32.37%), *Blastocystis* ambiguous taxon (25.49%), *D. fragilis* (12.12%), *P. hominis* (10.13%), and *Entamoeba* (3.45%).

Others have also used sequencing of 16S rRNA amplicon libraries to study the bacterial gut microbiome of people living in India [17–21, 42]. A study [17] of the gut microbiomes of 43 healthy individuals from urban and semi-urban districts in Maharashtra state (in the West of India, including Mumbai) also found that the microbial community at genus level was dominated by *Prevotella* (34.7%), *Bacteroides* (15.2%), and *Faecalibacterium* (5.6%), with additional prominant species *Megasphaera* (4.7%) and *Dialister* (3.9%). Similarly, the genera *Prevotella* (4.5%) and *Megasphaera* (8.5%) predominated in the gut microflora of 34 healthy Indian subjects from two urban cities (Delhi from the North; Pune from the West, also near Mumbai) [42] and were reported to be a distinctive feature of Indian gut microbiota compared with western cultures and to be a component of sharing OTUs with omnivorous mammals [42]. Elsewhere in India [21], the gut microbiome of a cohort (N=110) of healthy subjects from North-Central India (Bhopal City, Madhya Pradesh) consuming a primarily plant-based diet was dominated by *Prevotella*, whereas that of omnivorous individuals from Southern India (Kerala) showed more prominent associations with *Bacteroides*, *Ruminococcus* and *Faecalibacterium*. This aligned with previously described human gut microbiome enterotypes [16], with a high proportion of the North-Central population associated with enterotype-2 (73.5%) driven by *Prevotella* compared to the South India population sample (54% enterotype-2) where enterotype-1 (30.3%) driven by *Bacteroides* and enterotype-3 (16.1%) driven by *Ruminococcus* were also prevalent. This study also showed that species belonging to the genera *Bacteroides*, *Alistipes*, *Clostridium*, and *Ruminococcus* were relatively depleted in the Indian population compared to China, Denmark and the USA, whereas *Prevotella*, *Mitsuokella*, *Dialister*, *Megasphaera*, and *Lactobacillus* were the major drivers in separating Indian samples from other populations. In our study we found a predominance of enterotype-2 driven by *Prevotella*, with only a small number of individuals conforming to enterotype-1 driven by *Bacteroides* and a lack of enterotype-3 driven by *Ruminococcus*. High *Faecalibacterium* abundance was generally associated with high *Prevotella*. Bacterial taxa (e.g. *Dialister*, *Megasphaera*, *Mitsuokella*, *Lactobacillus*) previously shown to be characteristic of Indian gut microbiomes [17, 21, 42] occurred at <1% in our study population. Hence this population from North-East India does not fall within the criteria identified by these other studies to be characteristic of Indian populations.

One feature of the bacterial microbiota identified in our study was a high abundance of *Escherichia-Shigella* (9.1% aggregate abundance) in a subset of individuals that could define a different enterotype or sub-enterotype in our population compared to other Indian studies. This could reflect socio-economic status of residents of this region of India, together with higher prevalence of enteric diseases [43]. In a recent study of 475 patients with acute gastroenteritis Castaño-Rodriguez and colleagues [41] found that patient microbiomes clustered into three putative enterotypes, dominated by *Bacteroides*, *Escherichia-Shigella*, or *Faecalbacterium*. The novel *Escherichia-Shigella*-dominated enterotype found in a subset of patients was predicted to be more pro-inflammatory, and included the presence of *Enterobacteriaceae*, *Streptococcaceae*, and *Veillonellaceae*. In our study high *Escherichia-Shigella* abundance was associated with an overall dysbiosis of the gut bacterial flora, including a reduction in alpha diversity measures of species abundance and evenness. In our study we were also uniquely able to look at the interaction between prokaryotic and eukaryotic microflora in the gut. Interestingly, *Escherichia-Shigella* was also the major bacterial taxon driving the effect of eukaryotic *Blastocystis* abundance on bacterial beta diversity measures, with a negative correlation between *Blastocystis* and *Escherichia-Shigella* log_10_ counts. This might suggest that each out-competes the other and/or may reflect other influences of *Blastocystis* on gut microbial homeostasis (cf. below).

As alluded to above, our study was the first to employ 18S rRNA analysis of eukaryotic gut microflora in parallel with analysis of prokaryotic 16S rRNA sequences. Although overall measures of alpha and beta diversity for prokaryotic or eukaryotic microflora did not differ significantly between VL cases and EC, differences were observed at the level of individual bacterial or protozoan taxa. Specifically, using ANCOM we determined that EC had significantly higher levels of Ruminococcaceae UCG-014 (Family: Ruminococcaceae) and Gastranaerophilales_uncultured bacterium (Phylum: Cyanobacteria; Class: Melainabacteria; Order Gastranaerophilales) than VL cases. Both of these species are at relatively low aggregate relative abundance in our study population. However, previous studies in mice have shown that Ruminococcaceae UCG-014 is significantly reduced along with weight loss resulting from an intermittent fasting-mimicking diet [44]. This could reflect the effect of VL on cachexia and concomitant weight loss. Other research has noted perturbations of Ruminococcaceae UCG-014 in a murine model of inflammatory bowel disease [45] and in mice fed on low versus high salt diets [46]. The only published data on uncultured *Gastranaerophilales* spp. found an increase with aging in mice, which the authors suggested pointed to diminished antimicrobial defence with age [47]. Larger sample sizes and further research will be required to determine the functional significance, if any, of the reduced abundance of Ruminococcaceae UCG-014 and uncultured Gastranaerophilales in VL cases compared to EC.

A more prominent effect was the higher abundance of *P. hominis* in VL cases compared to EC. *P. hominis* is an anaerobic flagellated protist that colonizes the large intestine of a variety of mammals, including dogs, water buffalo, cattle, nonhuman primates, and humans [48, 49]. A recent epidemiological survey of faecal samples from northern China showed that 27% of dogs, 4% of human, and 47% of nonhuman primates carried *P. hominis* [48], with multiple authors suggesting that zoonotic transmission poses significant risk [48, 50, 51]. *P. hominis* has been associated with chronic diarrhoea and/or inflammatory bowel disease in animals [52], nonhuman primates [53] and humans [54], and is found in 14% of pregnant women in Papua New Guinea. In our study affected VL cases had higher abundance of *P. hominis* than EC. Due to their immuonological status VL cases may be more vulnerable to zoonotic transmission in the context of close association between humans, dogs and cattle/buffalo in this poor rural region of North-East India.

Independently of VL *versus* EC status, we observed high prevalence of *Blastocystis* spp. in our study population. Studies in a rural region of Brazil have also observed high prevalence of *Blastocystis*, with some evidence for genetic heterogeneity of *Blastocystis* subtypes within the population [55]. The two main genus-level taxa observed in our study, *Blastocystis* sp. MJ99-568 and *Blastocystis* ambiguous taxa, had similar effects on bacterial alpha and beta diversity measures. This included the effect of *Escherichia-Shigella* in driving the influence of *Blastocystis* on low bacterial beta-diversity, and the negative correlation between *Blastocystis* and *Escherichia-Shigella* counts as noted above. Using ANCOM we also determined that high *Blastocystis* was associated with low *Bacteroides* and with high *Clostridiales vadinBB60*. Given the former, the high prevalence of *Blastocystis* in our population could account for the low representation of bacterial enterotype-1 normally associated with *Bacteroides*. Within the gut *Bacteroides* are generally considered as friendly commensals [56]. Carbohydrate fermentation by *Bacteroides* produces volatile fatty acids that are absorbed through the large intestine and contribute to the host’s daily energy requirement [57]. High *Blastocystis* appears to put *Bacteroides* at a disadvantage thus influencing gut flora homeostasis. Another aspect of the dysbiosis associated with high *Blastocystis* is its association with high *Clostridiales vadinBB60*. Studies in mice have shown that abundance of *Clostridiales vadinBB60* is influenced by manipulation of meat protein in the diet and plays a part along with other microbiota in bidirectional signalling between gut and brain to regulate body metabolism [58]. However, nothing is known concerning pathogenicity of this species. In summary, whilst high *Blastocystis* was negatively correlated with *Escherichia-Shigella*, it likely contributes to overall dysbiosis of gut microflora in this North-East Indian population particularly to the low prevalence of enterotype-1 normally characterised by high *Bacteroides*.

Finally, an important aim of our study was to determine whether helminths could be detected using 18S rRNA analysis. This analysis yielded aggregate abundances of 0.78% for Ascaridida correlating with *Ascaris* identified microscopically, 0.77% for Rhabditida correlating with *Strongyloides* (hookworm) by microscopy, and 0.19% Cyclophyllidea correlating with *Hymenolepis* tapeworm by microscopy. The only apparent failure was detection of *Trichuris* using 18S rRNA in participants for whom small numbers of eggs had been identified in the stool samples, although we were able to identify Trichocephalida amplicons in one study participant negative by microscopy. Our inability to identify all Trichuris-infected individuals could also reflects loss of power in the 18S rRNA sample due to the presence of a large degree of contaminating bacterial 16S rRNA amplification caused by our demonstration that the EMP primers were not eukaryote-specific as originally reported. This has now been corrected on the EMP web-site. The use of eukaryote-specific 18S rRNA primers in future studies should allow more efficient detection of eukaryotic taxa in human faecal DNA, including the full range of soil transmitted helminth species. Thus, while the sample size employed in this scoping study was underpowered to determine any association between soil transmitted gut helminths and VL, our approach combining analysis of 16S rRNA and 18S rRNA analysis should work well in future studies designed to determine the interplay between gut microbiota and susceptibility to infectious diseases such as VL. Such studies may benefit initially by combining amplicon sequencing with a qPCR panel for eukaryotes which would also provide information on parasite load.

In conclusion, this scoping study has provided novel data on prokaryotic and eukaryotic gut microbiota for individuals living in this region of North-East India endemic for VL. Interesting dysbioses have been observed when considering the influence of putative pathogenic prokaryotic and eukaryotic species on global measures of bacterial diversity in the gut, and some important associations seen between VL and the eukaryotic pathogen *P. hominis*. Our study provides useful baseline data upon which to develop a broader analysis of pathogenic enteric microflora and their influence on gut microbial health and NTDs generally.

## Supporting Information Legends

**S1 Checklist. STrengthening the Reporting of OBservational studies in Epidemiology (STROBE) checklist.** Provides details of how our study complies with guidelines for epidemiological case-control studies set down by the STROBE consortium of international epidemiologists, methodologists, statisticians, researchers and journal editors.

S1 Table. Metadata for endemic control (EC) and visceral leishmaniasis (VL) individuals contributing to the study.

S2 Table. Genus-level raw counts for 16S rRNA analysis.

S3 Table. Genus-level raw counts for 18S rRNA analysis after filtering out a small number of human 18S rRNA sequences and the cross-contaminating bacterial 16S rRNA sequences.

**S1 Figure.** Rarefaction plots based on Shannon’s phylogenetic alpha diversity index for (a) 16S rRNA sequence date and (b) 18S rRNA sequence data (after filtering out contaminating bacterial sequences).

**S2 Figure. Bar plots for relative abundance of taxa for 18S rRNA data prior to filtering out contaminating bacterial species identified as unclassified “Eukaryota”.** Taxa ordered by relative abundance of the contaminant “Eukaryota”. (a) VL case and EC samples. (b) Negative (MD labels) and positive (MOCK labels) controls. Negative controls indicate a small degree of laboratory cross-contamination of taxa identified in experimental samples. (c) Bar plot for 16S taxa that were contaminants in the negative control samples, colour key to relative abundances as for main figure 1(a). Note that these percentages are relative to the much smaller number of processed reads for negative controls (9573±9294 compared to 71,123±22,122 for positive mock controls and 91,923±69,706 for VL cases and EC samples). The diversity within the samples was equivalent to VL cases and EC, indicating that they were likely due to cross-contamination in the endemic laboratory. The absolute numbers of reads were unlikely to have influenced the data and conclusions drawn for experimental samples.

**S3 Figure. Influence of age and gender on 16S rRNA-determined prokaryotic microbial profiles.** (a) and (b) bar plots for relative abundance of taxa by age and sex, respectively; colour key to relative abundances as for main figure 1(a). (c) to (f) comparisons of species richness (number of ASVs) and alpha diversity measures (as labelled). (g) PCoA plots (left to right: PC1xPC2; PC1xPC3; PC2xPC3) for weighted UniFrac beta diversity by gender (see key). (h) Three-dimensional PCoA plot for weighted UniFrac beta diversity by age (see key for continuous scale).

**S4 Figure. Influence of age and gender on 18S rRNA-determined eukaryotic microbial profiles.** (a) and (b) bar plots for relative abundance of taxa by age and sex, respectively; colour key to relative abundances as for main figure 1(a). (c) to (f) comparisons of species richness (number of ASVs) and alpha diversity measures (as labelled). (g) PCoA plots (left to right: PC1xPC2; PC1xPC3; PC2xPC3) for weighted UniFrac beta diversity by gender (see key). (h) Three-dimensional PCoA plot for weighted UniFrac beta diversity by age (see key for continuous scale).

**S5 Figure. Influence of micro-geography on 16S rRNA-determined prokaryotic microbial profiles.** (a) map of the district of Muzaffarpur showing subdistricts or block, annotated with the numbers of samples collected from different blocks. Listed below the map are numbers of samples from blocks outside of Muzaffarpur. (b) bar plots for relative abundance of taxa by district; colour key to relative abundances of taxa as for main figure 1(a). (c) to (f) comparisons of species richness (number of ASVs) and alpha diversity measures (as labelled) by blocks with N≥4. (g) PCoA plots (left to right: PC1xPC2; PC1xPC3; PC2xPC3) for weighted UniFrac beta diversity by colour-coded blocks.

**S6 Figure. Influence of micro-geography on 18S rRNA-determined prokaryotic microbial profiles.** (a) map of the district of Muzaffarpur showing subdistricts or block, annotated with the numbers of samples collected from different blocks. Listed below the map are numbers of samples from blocks outside of Muzaffarpur. (b) bar plots for relative abundance of taxa by district; colour key to relative abundances of taxa as for main figure 2(a). (c) to (f) comparisons of species richness (number of ASVs) and alpha diversity measures (as labelled) by blocks with N≥3. (g) PCoA plots (left to right: PC1xPC2; PC1xPC3; PC2xPC3) for Bray’s dissimilarity beta diversity by colour-coded blocks.

## Acknowledgements

We would like to thank the hospital staffs at Kala–azar Medical Research Centre, Muzaffarpur for their assistance in the collection of samples and all research scholars of Infectious Disease Research Laboratory, Banaras Hindu University for their kind help during the study.

## Author Contributions

**Conceptualization:** Rachael Lappan, Cajsa Classon, Om-Prakash Singh, Jenefer M. Blackwell

**Data curation:** Rachael Lappan, Cajsa Classon, Om-Prakash Singh,

**Formal analysis:** Rachael Lappan, Ricardo de Almeida, Jenefer M. Blackwell

**Funding acquisition:** Shyam Sundar, Jenefer M. Blackwell

**Investigation:** Rachael Lappan, Cajsa Classon, Shashi Kumar, Om-Prakash Singh, Poonam Kumari, Sangeeta Kansal, Jaya Chakravarty

**Project administration:** Cajsa Classon, Om-Prakash Singh, Sangeeta Kansal, Jaya Chakravarty

**Supervision:** Shyam Sundar, Jenefer M. Blackwell

**Validation:** Rachael Lappan, Cajsa Classon, Jenefer M. Blackwell

**Visualization:** Rachael Lappan, Jenefer M. Blackwell

**Writing – original draft:** Jenefer M. Blackwell

**Writing – reviewing and editing:** Jenefer M. Blackwell, Rachael Lappan, Cajsa Classon, Om-Prakash Singh.

**Approval of manuscript:** All authors read and approved the manuscript.

## Data Availability Statement

The dataset generated and analysed during the current study is available in the NCBI Sequence Read Archive (PRJNA525566; available at https://www.ncbi.nlm.nih.gov/sra/PRJNA525566). Complete documentation of all 16S rRNA and 18S rRNA sequence analysis code can be found at https://rachaellappan.github.io/VL-QIIME2-analysis/.

